# Multiple mechanisms of action of an extremely painful venom

**DOI:** 10.1101/2024.09.12.612741

**Authors:** Lydia J. Borjon, Luana C. de Assis Ferreira, Jonathan C. Trinidad, Sunčica Šašić, Andrea G. Hohmann, W. Daniel Tracey

**Author notes:** Corresponding author and lead contact: W. Daniel Tracey.

## Abstract

Evolutionary arms races between predator and prey can lead to extremely specific and effective defense mechanisms. Such defenses include venoms that deter predators by targeting nociceptive (pain-sensing) pathways. Through co-evolution, venom toxins can become extremely efficient modulators of their molecular targets. The venom of velvet ants (Hymenoptera: Mutillidae) is notoriously painful. The intensity of a velvet ant sting has been described as “Explosive and long lasting, you sound insane as you scream. Hot oil from the deep fryer spilling over your entire hand.” [1] The effectiveness of the velvet ant sting as a deterrent against potential predators has been shown across vertebrate orders, including mammals, amphibians, reptiles, and birds [2–4]. The venom’s low toxicity suggests it has a targeted effect on nociceptive sensory mechanisms [5]. This leads to the hypothesis that velvet ant venom targets a conserved nociception mechanism, which we sought to uncover using *Drosophila melanogaster* as a model system. *Drosophila* larvae have peripheral sensory neurons that sense potentially damaging (noxious) stimuli such as high temperature, harsh mechanical touch, and noxious chemicals [6–9]. These polymodal nociceptors are called class IV multidendritic dendritic arborizing (cIV da) neurons, and they share many features with vertebrate nociceptors, including conserved sensory receptor channels [10,11]. We found that velvet ant venom strongly activated *Drosophila* nociceptors through heteromeric Pickpocket/Balboa (Ppk/Bba) ion channels. Furthermore, we found a single venom peptide (Do6a) that activated larval nociceptors at nanomolar concentrations through Ppk/Bba. *Drosophila* Ppk/Bba is homologous to mammalian Acid Sensing Ion Channels (ASICs) [12]. However, the Do6a peptide did not produce behavioral signs of nociception in mice, which was instead triggered by other non-specific, less potent, peptides within the venom. This suggests that Do6a is an insect-specific venom component that potently activates insect nociceptors. Consistent with this, we showed that the velvet ant’s defensive sting produced aversive behavior in a predatory praying mantis. Together, our results indicate that velvet ant venom evolved to target nociceptive systems of both vertebrates and invertebrates, but through different molecular mechanisms.

## Results and Discussion

### Velvet ant venom activates larval nociceptors

We first tested whether velvet ant venom activates *Drosophila* nociceptors. Venom samples were collected from female Scarlet Velvet Ants (*Dasymutilla occidentalis*, Fig. 1A) by inducing them to sting into a piece of parafilm and deposit venom droplets. Larval peripheral sensory neurons were exposed in a semi-intact fillet preparation [13], making it possible to apply venom directly to neurons whose dendritic arbors remain intact and embedded in their native tissue and cellular milieu. Sensory neuron activity was assessed with cell-type specific expression of the genetically encoded calcium sensor GCaMP6f [14], and fluorescent time-series images were obtained on a high-speed confocal microscope. Application of dilute venom (estimated dilution factor of more than 1:20,000) activated only nociceptive cIV da neurons (Fig. 1B-C, Video S1). Other sensory neurons, such as class III dendritic arborizing (cIII da) neurons, which are mainly responsible for sensing innocuous mechanical stimuli and play a contributory role in larval nociception [7,15,16], did not respond to diluted venom applications (Fig. 1B-C, Video S1). These observations demonstrate that velvet ant venom activates the nociceptors of an insect.

**Figure 1.**
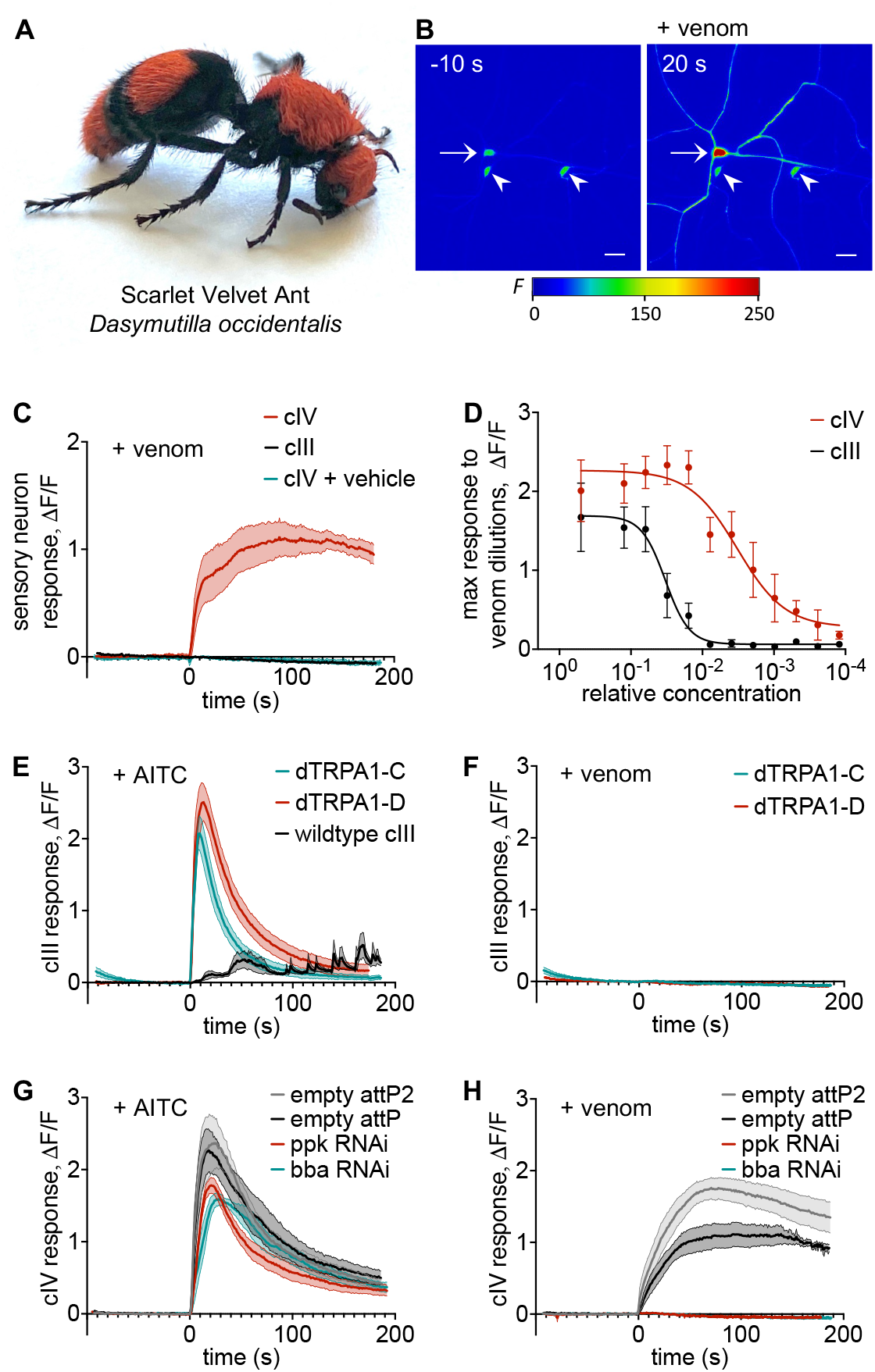
Velvet ant venom activates larval sensory neurons with a potent nociceptor-specific component that requires Pickpocket/Balboa. (**A**)Venom was obtained from collected individuals of the female Scarlet Velvet Ant (*Dasymutilla occidentalis)*. (**B**)Still images from time-series optical recording of cIV da (arrow) and cIII da (arrow heads) neurons expressing the genetically encoded Ca^2+^ sensor GCaMP6f (*w; ppk1*.*9-Gal4 UAS-GCaMP6f*) 10 s before and 20 s after application of diluted venom. Scale bar represents 20μm. Pseudo-coloring for fluorescence intensity (**C**). Calcium imaging of diluted venom or vehicle application at time 0 s. cIV da neurons (*w; ppk1*.*9-Gal4 UAS-GCaMP6f/+*) respond to venom but not vehicle application. cIII da neurons (*w; nompC-Gal4/+; UAS-GCaMP6f/+*) do not respond to venom application. n = 6-12 neurons. (**D**)Dose-response to serial dilutions of pooled venom samples in cIII da and cIV da neurons (*w; ppk1*.*9-Gal4 UAS-GCaMP6f*). A nociceptor-specific component potently activates cIV da neurons at >100 fold greater dilution relative to venom that activates cIII da neurons. Data represented as mean ± SEM fit with a Sigmoid curve, at each dilution n = 1-14 neurons from 3 venom pools. See also Fig. S1. (**E**)Calcium imaging of AITC application at time 0 s. cIII da neurons (*w; nompC-Gal4/+; UAS-GCaMP6f/+*) do not activate strongly in response to AITC. Expression of *dTRPA1-C* or *dTRPA1-D* in cIII da neurons (*w; nompC-Gal4/UAS-dTRPA1-C; UAS-GCaMP6f/+* or *w; nompC-Gal4/UAS-dTRPA1-D; UAS-GCaMP6f/+*) renders them responsive to AITC. n = 12 neurons. (**F**)Calcium imaging of venom application at time 0 s. Expression of *dTRPA1-C* or *dTRPA1-D* in cIII da neurons (*w; nompC-Gal4/UAS-dTRPA1-C; UAS-GCaMP6f/+* or *w; nompC-Gal4/UAS-dTRPA1-D; UAS-GCaMP6f/+*) does not render them responsive to venom. n = 12 neurons. (**G**)Calcium imaging of AITC application at time 0 s. cIV da neurons with RNAi against *ppk* (*w; ppk1*.*9-Gal4 UAS-GCaMP6f/UAS-ppk RNAi; UAS-dicer2/+*) or against *bba* (*w; ppk1*.*9-Gal4 UAS-GCaMP6f/UAS-bba RNAi; UAS-dicer2/+*) respond to AITC similarly as genetic controls (*w; ppk1*.*9-Gal4 UAS-GCaMP6f/attP; UAS-dicer2/+* or *w; ppk1*.*9-Gal4 UAS-GCaMP6f/attP2; UAS-dicer2/+*), showing that knock-down of *ppk* or *bba* does not interfere with the neurons’ overall ability to respond to stimuli. n = 6 neurons. (**H**)Calcium imaging of venom application at time 0 s. RNAi against *ppk* (*w; ppk1*.*9-Gal4 UAS-GCaMP6f/UAS-ppk RNAi; UAS-dicer2/+*) or against *bba* (*w; ppk1*.*9-Gal4 UAS-GCaMP6f/ UAS-bba RNAi; UAS-dicer2/+*) eliminates the response to venom in cIV da neurons, while genetic controls (*w; ppk1*.*9-Gal4 UAS-GCaMP6f/attP; UAS-dicer2/+* or *w; ppk1*.*9-Gal4 UAS-GCaMP6f/attP2; UAS-dicer2/+*) respond normally to venom. n = 6-7 neurons. C, E-H: Data represented as mean ± SEM. See also Video S1.

Although cIII da neurons were not activated by dilute venom, we found that more concentrated samples of whole venom pooled from multiple velvet ants could evoke a broader activation in all classes of da sensory neurons. Testing serial dilutions of the concentrated venom samples on cIV da and cIII da neurons simultaneously, revealed that high concentrations activate both neuron classes (Fig 1D, Fig. S1A-C, Video S1). However, cIII da responses disappeared after dilutions of 1:128 relative to the starting concentration of the pooled venom samples (estimated dilution factor of ∼1:1,000 relative to pure venom). The cIV da nociceptor-specific responses were still observed for pooled venom samples diluted to 1:8192 (estimated dilution factor of ∼1:80,000 relative to pure venom), almost 100 times more dilute than those that activated cIII da. This indicates that the most potent venom activity acts specifically on larval nociceptors. Thus, we hypothesized that a component of the venom targets a molecule that is found in the cIV da nociceptors but not in other da sensory neurons. We next sought to identify this potent active component and its molecular target.

### Velvet ant venom does not activate nociceptor-specific dTRPA1 channels

We expected the molecular target of velvet ant venom to be a sensory receptor channel whose expression is specific to cIV da neurons. We first chose to investigate *Drosophila* TRPA1 (dTRPA1) as a likely candidate target because it is highly conserved across animals, and it is involved in the sensation of noxious temperature as well as a wide array of irritant compounds [17–22], including allyl isothiocyanate (AITC), the pungent component of mustard and wasabi [23–25]. Given TRPA1’s versatility in detecting many compounds with different chemical properties, it seemed like an ideal target for the evolution of a venom that could act as a deterrent against organisms across the animal kingdom.

dTRPA1 has five alternative splice isoforms in flies [26–28], and two of them (dTRPA1-C and dTRPA1-D) are expressed specifically in cIV da neurons in the larval peripheral nervous system [26,28]. Ectopic expression of either dTRPA1-C or dTRPA1-D isoforms in cIII da neurons, which normally do not respond either to the TRPA1 agonist AITC or to venom, rendered them sensitive to AITC (Fig. 1E, Video S1), but not to venom (Fig. 1F, Video S1). Thus, dTRPA1 was not likely to be a venom target.

### Pickpocket/Balboa channels are necessary for nociceptor responses to velvet ant venom

Next, we investigated Pickpocket/Balboa (Ppk/Bba, also known as Ppk1/Ppk26), another candidate target channel whose expression is restricted to cIV da neurons in the larval peripheral nervous system. Ppk/Bba is required for mechanical nociception in fly larvae [29–31], but this channel has not previously been shown to be activated by ligands. Ppk/Bba is homologous to DEG/ENaC and ASIC channels which comprise a large channel family with polymodal sensitivity to noxious stimuli including low pH, mechanical stimuli, and inflammatory ligands [32–36]. Furthermore, mammalian ASIC’s are targets of multiple venoms from other animals [37–41]. Ppk and Bba form heteromeric channels, and the absence of either subunit disrupts proper channel localization in the plasma membrane [29,31,42].

RNAi knock-down of either *ppk* or *bba* in cIV da neurons did not strongly affect their responses to AITC (Fig 1G, Video S1), showing that nociceptors lacking these channels retain their excitability in response to this noxious chemical stimulus. However, consistent with the hypothesis of Ppk/Bba as a venom target, RNAi knock-down of either *ppk* or *bba* in cIV da neurons eliminated their response to venom (Fig. 1H, Video S1). This result suggests that the venom of the Scarlet Velvet Ant has a specific channel target that results in nociceptor-specific activation in an insect.

### Co-expression of Pickpocket and Balboa is sufficient for venom sensitivity in neurons

To further test whether Ppk/Bba is a venom target, we ectopically expressed *ppk* and *bba* in cIII da neurons, which normally do not respond to diluted venom. Expression of *ppk* alone (Fig. 2A-B, Video S2) or *bba* alone (Fig. 2C-D, Video S2) did not render cIII da neurons sensitive to venom. This is consistent with the expectation that both channel subunits need to be present for proper subcellular localization [29,31,42]. Remarkably, co-expression of *ppk* and *bba* rendered cIII da neurons sensitive to venom (Fig. 2E-F, Video S2), but not to application of vehicle (Fig. 2G-H, Video S2). This observation, combined with loss of function experiments above, demonstrates that Ppk/Bba is a molecular target of velvet ant venom. Our data also provide the first functional demonstration that co-expression of Ppk and Bba results in functional channels that lead to neuronal activation.

**Figure 2.**
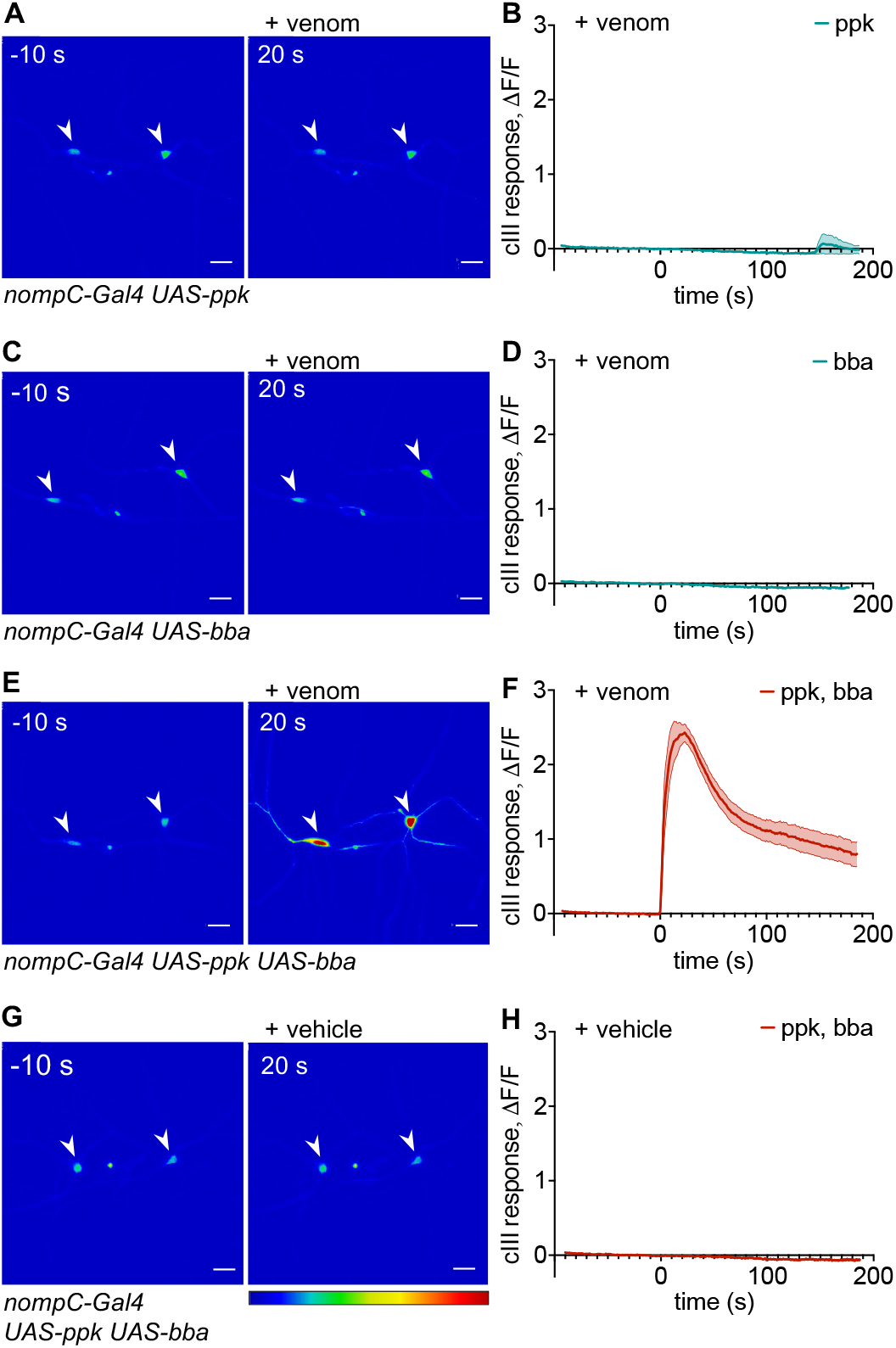
Pickpocket/Balboa channels are sufficient for neuronal response to venom. **(A-B)** Still images (A) and quantification (B) of diluted venom application at time 0 s on cIII da neurons expressing *ppk* alone (*w; nompC-Gal4/+; UAS-GCaMP6f/UAS-ppk*), which do not respond to venom. n = 12 neurons. **(C-D)** Still images (C) and quantification (D) of diluted venom application at time 0 s on cIII da neurons expressing *bba* alone (*w; nompC-Gal4/+; UAS-GCaMP6f/UAS-bba-mCherry*), which do not respond to venom. n = 12 neurons. **(E-F)** Still images (E) and quantification (F) of diluted venom application at time 0 s on cIII da neurons co-expressiing *ppk* and *bba* (*w; nompC-Gal4/+; UAS-GCaMP6f/UAS-ppk UAS-bba-mCherry*), which respond to venom. n = 16 neurons. **(G-H)** Still images (G) and quantification (H) of vehicle application at time 0 s on cIII da neurons co-expressing *ppk* and *bba* (*w; nompC-Gal4/+; UAS-GCaMP6f/UAS-ppk UAS-bba-mCherry*), which do not respond to vehicle. n = 12 neurons. B, D, F, H: Data represented as mean ± SEM. See also Video S2.

### A single venom peptide with potent nociceptor specific effects targets Pickpocket/Balboa

We next sought to identify the venom component responsible for the nociceptor-specific activation. The venom of *Dasymutilla occidentalis* is composed primarily of short peptides (7-68 aa) whose sequences were determined by Jensen et al. (2021) using a combined transcriptomic and HPLC/mass spectrometry approach [43]. We chemically synthesized each of the 24 peptides and screened them on larval nociceptive neurons (except for 2 peptides (Do14a, Do17a) which we were not able to test due to insolubility). Each peptide was initially tested at 100 μM, revealing 4 peptides (Do6a, Do10a, Do12a, Do13a) that activated larval sensory neurons (Fig. 3A-B, Video S3).

**Figure 3.**
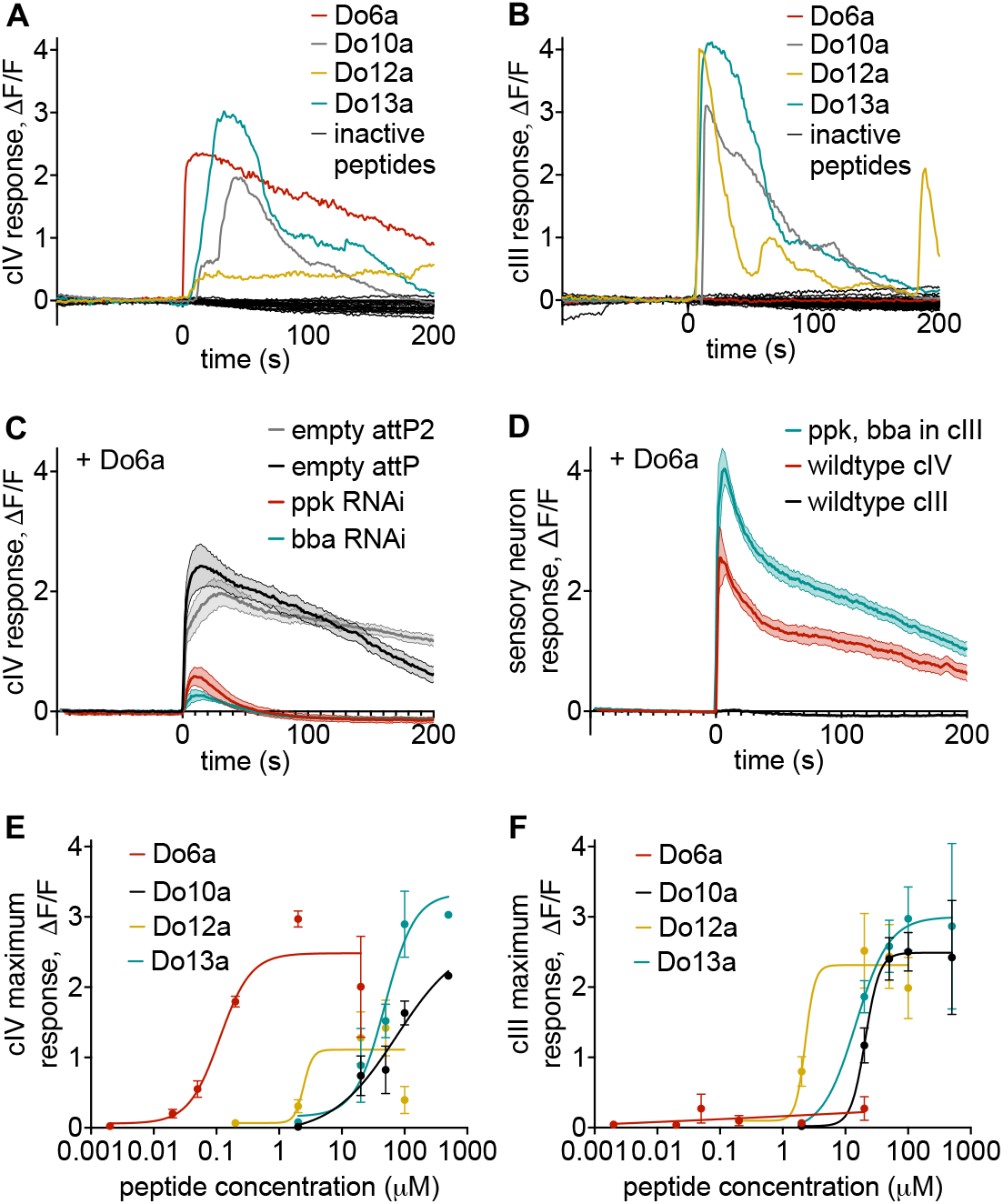
Four venom peptides activate larval sensory neurons but only one peptide, Do6a, mediates activation through Pickpocket/Balboa. **(A)**Exemplar calcium-imaging traces in cIV da neurons (*w; ppk1*.*9-Gal4 UAS-GCaMP6f*) in response to all venom peptides tested at 100 μM and applied at time 0 s. Four peptides, Do6a, Do10a, Do12a, and Do13a, activate cIV da neurons. The eighteen other peptides do not activate larval sensory neurons. Two peptides, Do14a and Do17a, were not tested due to poor solubility. (**B**)Exemplar calcium-imaging traces in cIII da neurons (*w; ppk1*.*9-Gal4 UAS-GCaMP6f*) in response to venom peptides tested at 100 μM and applied at time 0 s (from the same trials shown in (A)). Peptides Do10a, Do12a, and Do13a activate cIII da neurons, but peptide Do6a does not. The remaining peptides also do not activate cIII da neurons. (**C**)Calcium imaging of Do6a (20 μM) application at time 0 s. RNAi against *ppk* (*w; ppk1*.*9-Gal4 UAS-GCaMP6f/UAS-ppk RNAi; UAS-dicer2/+*) or against *bba* (*w; ppk1*.*9-Gal4 UAS-GCaMP6f/ UAS-bba RNAi; UAS-dicer2/+*) eliminates the response to Do6a in cIV da neurons, while genetic controls (*w; ppk1*.*9-Gal4 UAS-GCaMP6f/attP; UAS-dicer2/+* or *w; ppk1*.*9-Gal4 UAS-GCaMP6f/attP2; UAS-dicer2/+*) respond normally to Do6a. Data represented as mean ± SEM, n = 6 neurons. (**D**)Calcium imaging of Do6a (20 μM) application at time 0 s. cIV da neurons (*w; ppk1*.*9-Gal4 UAS-GCaMP6f/+*) respond to Do6a application but cIII da neurons (*w; nompC-Gal4/+; UAS-GCaMP6f/+*) do not. Co-expression of *ppk* and *bba* in cIII da neurons (*w; nompC-Gal4/+; UAS-GCaMP6f/UAS-ppk UAS-bba-mCherry*) renders them responsive to Do6a. Data represented as mean ± SEM, n = 6-12 neurons. See also Fig. S2. (**E**)Dose-response curves in cIV da neurons (*w; ppk1*.*9-Gal4 UAS-GCaMP6f*) for Do6a, Do10a, Do12a, and Do13a. Half-maximal effective concentration (EC_50_) is 113 nM for Do6a, 74.5 μM for Do10a, 2.45 μM for Do12a, and 48.7 μM for Do13a. Data represented as mean ± SEM fit with a Sigmoid curve, n = 3 neurons per concentration. (**F**) Dose-response curves in cIII da neurons (*w; ppk1*.*9-Gal4 UAS-GCaMP6f*) for Do6a, Do10a, Do12a, and Do13a (from the same trials as in (E)). Half-maximal effective concentration (EC_50_) is 20.7 μM for Do10a, 2.29 μM for Do12a, and 14.7 μM for Do13a. Data represented as mean ± SEM fit with a Sigmoid curve, n = 3-10 neurons per concentration. See also Video S3.

Only a single peptide, Do6a, specifically activated nociceptive cIV da neurons and not cIII da neurons (Fig. 3A-B, Video S3). The other 3 peptides (Do10a, Do12a, Do13a) also activated cIV da nociceptors, but at higher concentrations and at slower time scales than Do6a. In addition, these 3 peptides have a general effect that also activates cIII da sensory neurons (Fig. 3A-B, Video S3).

Furthermore, the nociceptor-specific activity of Do6a is mediated through Ppk/Bba channels as demonstrated by both RNAi and ectopic expression experiments. RNAi against either *ppk* or *bba* in nociceptors nearly eliminated the response to Do6a, recapitulating the result seen with whole venom (Fig. 3C, Video S3). In addition, co-expression of *ppk* and *bba* rendered cIII da neurons sensitive to Do6a (Fig. 3D, Video S3). This was also the case in class I da proprioceptive neurons (Fig. S2), the cell type in which it was previously shown that ectopic expression of both subunits leads to proper sub-cellular localization [29]. These observations collectively suggest that a single peptide targeting heteromeric Ppk/Bba channels is responsible for the potent nociceptor-specific activity of velvet ant venom.

The dose-response curve for Do6a showed a half-maximal effective concentration (EC_50_) of 113 nM (Fig. 3E). Meanwhile, the EC_50_’s of Do10a, Do12a, and Do13a were in micromolar ranges for their activity in both cIV da and cIII da neurons (75 μM, 2.5 μM, 49 μM respectively in cIV da, and 21 μM, 2.3 μM, 15 μM respectively in cIII da, Fig. 3E-F). However, the contribution of each peptide to venom function also depends on its abundance in the whole venom cocktail. To investigate this relationship, we performed LC-MS to quantify peptide concentrations by comparing the peaks from whole venom to standard mixtures of synthetic peptides at known concentrations. We analyzed two venom samples that were pooled from multiple velvet ants, and we were able to identify 21 of the expected peptides (Do8a, Do9a, and Do14a were not detected, Supplemental Table S1). Some peptide quantities varied considerably between the two samples, suggesting that venom contents may vary between individuals, between stings, or across time. However, the relative amounts of the neuron-activating peptides Do6a, Do10a, Do12a, and Do13a were similar in both samples. Do6a was the most abundant peptide in the venom and was 1-3 orders of magnitude more abundant than Do10a, Do12a, and Do13a. Thus, Do6a is both more potent and more concentrated in whole venom than the peptides with non-specific neuronal effects. We therefore propose that Do6a is the major component of the Scarlet Velvet Ant venom that functions to activate insect nociceptors.

### Venom components are noxious to mice, but do not involve Do6a

After identifying Do6a as an important noxious venom component targeting insects, we asked whether the same molecules are responsible for the painful effects of venom in vertebrates.

Mammalian nociceptors express ASIC ion channels that are homologous to the Do6a target Ppk/Bba, and thus represent a likely avenue for a conserved mechanism for this widely effective venom. We confirmed that velvet ant venom is noxious to mice by injecting venom samples unilaterally into mouse hind paws. Compared to a control vehicle injection, the mice injected with venom showed pronounced nocifensive behaviors, such as licking, shaking, or tapping the affected paw during 10 minutes post-injection (Fig. 4A). They also exhibited robust mechanical hypersensitivity in the ipsilateral (venom injected) paw (Fig. 4B) but not in the contralateral (intact) paw (Fig. 4C) for 30 min post-injection.

**Figure 4.**
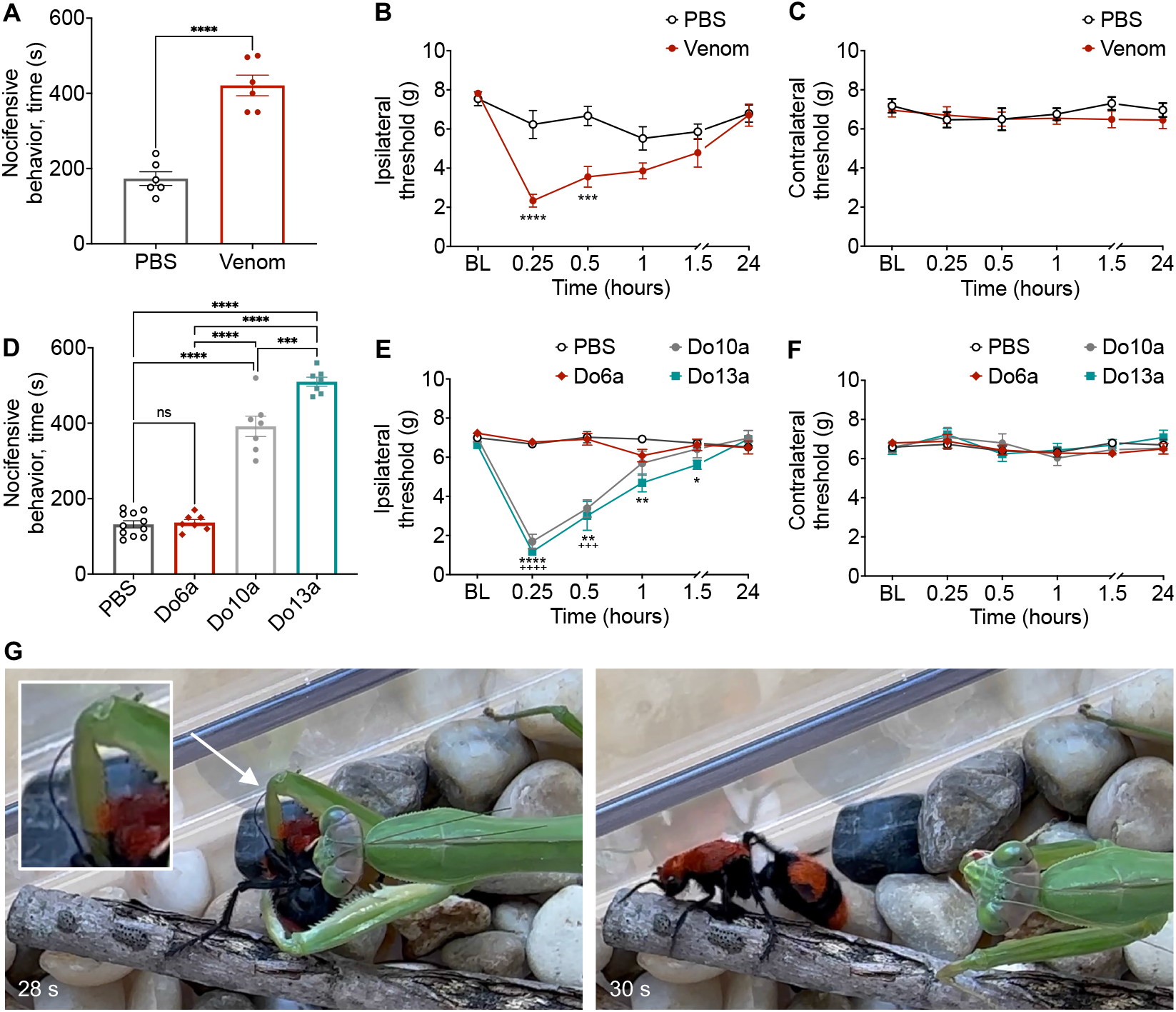
Whole venom and venom peptides are noxious to mice, and velvet ant stings deter an insect predator. (**A**)Intraplantar injection of venom unilaterally into the mouse hind paw produced nocifensive behaviors compared to control PBS injection. Nocifensive behaviors were measured over 10 min following injection of venom or PBS into the hind paw. ****p<0.0001, unpaired two-tailed t-test. (**B**)Venom decreases paw withdrawal thresholds (force in g) in the ipsilateral (injected) paw compared to PBS injection. Withdrawal thresholds were measured at the pre-injection baseline (BL) and at various time points after the intraplantar injection. Two-way repeated measures ANOVA with main effect between groups F(1, 10) = 15.57, p = 0.0027, Sidak’s multiple comparison test, ****p<0.0001, ***p<0.001. (**C**)Intraplantar injections (same mice shown in B) did not alter paw withdrawal thresholds in the contralateral (intact) paw. Two-way repeated measures ANOVA with no main effect between groups F(1, 10) = 0.6405, p = 0.4421, Sidak’s multiple comparison test with no significant differences. (**D**)Intraplantar injection of peptides Do10a and Do13a but not Do6a produced nocifensive behavior in the injected paw. Do13a produced maximal nocifensive behavior compared to all other groups. Two-way ANOVA with main effect between groups F(3, 18) = 121.7, p < 0.0001, Tukey’s multiple comparison test, ****p<0.0001, ***p<0.001, ns no significant difference. (**E**)Intraplantar injection of peptides Do10a and Do13a but not Do6a decreased mechanical paw withdrawal thresholds in the injected paw. Two-way repeated measures ANOVA with main effect between groups F(3, 28) = 42.94, p < 0.0001, Tukey’s multiple comparison test, ****p<0.0001, ***p<0.001, **p<0.01, *p<0.05. + denotes differences between PBS and Do10a, * denotes differences between PBS and Do13a. (**F**)None of the treatments altered paw withdrawal thresholds in the contralateral (intact) paw (same mice shown in E). Two-way repeated measures ANOVA with no main effect between groups F(3, 28) = 0.1698, p = 0.9159, Tukey’s multiple comparison test with no significant differences. A-F: Data represented as mean ± SEM, n = 6-11 mice per group. (**G**)Still frames from Video S4 showing the defensive sting inflicted by the velvet ant to the foreleg femur/trochanter of the praying mantis, followed by the velvet ant’s escape a few seconds later.

We next tested whether the same peptides that activate sensory neurons in fly larvae are also noxious to mice. Peptides Do6a, Do10a, and Do13a were each individually injected into the hind paw of separate groups of mice under blinded conditions and the mice were assessed for nocifensive behaviors and subsequent mechanical hypersensitivity. Intraplantar injection of Do10a and Do13a each produced marked increases in nocifensive behaviors (Fig. 4D), as well as mechanical hypersensitivity lasting more than 60 min post-injection (Fig. 4E). Surprisingly, effects of Do6a injection were indistinguishable from control injection (Fig. 4D-E). None of the peptides altered paw withdrawal thresholds in the intact paw contralateral to the injection (Fig. 4F). While two venom peptides (Do10a and Do13a) are noxious to mice, and these peptides can also activate insect neurons, the most potent peptide for insect nociceptors (Do6a) did not produce a nocifensive response or apparent behavioral hypersensitivity in mice, even at concentrations that are ∼10,000x higher than the EC_50_ for larval nociceptor activation. Therefore, the nociceptive mechanism of action of the venom is different in mammals and insects.

Consistent with our observations, a previous study showed that venom peptides from other velvet ant species that are homologous to Do10a and Do13a (Dk13a and Dk5a from *Dasymutilla klugii*, and Dg3a and Dg6a from *Dasymutilla gloriosa*, Fig. S3) activate dissociated mouse sensory neurons in a non-specific manner [43]. The authors proposed that the non-specific activation of multiple cell types by these peptides is mediated by a pore-forming mechanism [43]. This may be consistent with our observation that the effect of the non-specific peptides on larval sensory neurons was variable and often delayed, which suggests a mechanism that is not mediated by gating of a receptor ion channel. Although *D. klugii* venom also contains peptides with amino acid sequence homology to Do12a (Dk9a and Dk9b, Fig. S3), these peptides were not reported to activate mouse neurons [43]. Remarkably, each of the velvet ant species investigated had a venom peptide that is nearly identical to Do6a (Fig. S3), but its potential importance as a defensive substance was only revealed through the study of insect neurons.

### Velvet ant stings are a defense against an insect predator

Do6a activates nociceptive neurons and nociceptive receptor channels that are known to respond to other noxious stimuli in *Drosophila*, but it did not produce apparent pain behaviors in mice. We therefore wondered what evolutionary pressures would lead to this potent, apparently insect-specific venom component. We hypothesized that, in addition to vertebrate predators against whom velvet ant defenses have previously been shown to be effective [2–4], they may also have insect predators that could be deterred by the venomous sting. Given the diversity of insects, velvet ants in the wild are likely to encounter a multitude of predatory insects. Further, homologous *pickpocket* genes are prevalent across insects [44–46]. We asked, therefore, whether velvet ant venom triggers nocifensive behaviors in insects.

Mantises (Mantodea: Mantidae), which are a widespread group with over 2500 species [47], are among potential predators of velvet ants. Indeed, we found that when presented with a velvet ant, a praying mantis (female *Tenodera sinensis*) readily attacked and attempted to consume it. Yet, despite the mantis’s extremely strong grip, the velvet ant defended itself by deploying its stinger in multiple attempts to sting the mantis. In 5 out of 5 trials the velvet ant escaped when the mantis startled and suddenly released it. We clearly documented one instance when the mantis released the prey at the moment that the stinger penetrated the mantis foreleg femur/trochanter (Fig. 4G, Video S4). We also observed and documented that the release of the velvet ant was sometimes followed by grooming of the site of the sting and apparent discoordination of the mantis foreleg for several minutes, actions which resemble established nocifensive behaviors in other species (Video S4). The mantis always survived these attacks which differs from a previously reported lethal effect of velvet ant venom on insects [43]. These observations suggest that the venom serves the same function when acting on insects and on vertebrates, which is to deter potential predators by activating their nociceptive system. The molecular details of how this activation occurs, however, is different in insects and mammals.

## Conclusion

Our study has several unexpected findings: (1) velvet ant venom, which is extremely effective at triggering pain in humans, also activates the nociceptive neurons of an insect; (2) a single venom peptide (Do6a) is responsible for potent activation of insect nociceptors and is also the most abundant peptide within the venom; (3) Do6a does not produce signs of nociception in mice, indicating that it is an insect-specific venom mechanism; (4) velvet ant stings are sufficiently aversive to override the prey drive of a predatory insect.

Because the venom can activate neurons of such distant phyla as insects and mammals we expected to find that it acts through an evolutionarily ancient nociceptive mechanism. Instead, we uncovered that the venom cocktail has multiple mechanisms: a general mechanism that is widely effective across the animal kingdom, and a specific mechanism that is tailored to target the nociceptive system of insects. That the most abundant venom peptide acts specifically against insects suggests that interactions with predatory insects is a more important selective pressure for the evolution of velvet ant venom than interactions with vertebrates. Velvet ants are parasitoids of ground dwelling bees and wasps [48] and it is possible that the venom is also used in defensive encounters that may happen when they enter the nests of these hosts.

Despite their ability to avoid potentially dangerous stimuli, whether insects “feel pain” remains a subject of considerable debate and controversy [49,50]. Our results show that, similar to defensive compounds that evoke noxious sensations by activating pain receptors in mammals (eg. capsaicin of pepper plants), insects can exploit nociceptive pathways of other insects to trigger aversion. Although the mechanism of action for velvet ant venom is distinct in mammals and insects, the fact that the venom converged to target the analogous sensory system in these widely divergent taxa provides further support for the functional similarity between insect nociception and mammalian pain systems.

## Acknowledgments

The authors would like to thank community members who assisted with velvet ant collection. We thank the Yuh-Nung Jan lab for providing *UAS-bba::mCherry* flies, and the Bloomington Drosophila Stock Center (NIH P40OD018537) for providing fly lines and other fly resources. We thank Jeremy Borjon and Elizabeth Haswell for careful reading and comments on earlier manuscript drafts. This study was supported by grants from the National Institute of Health 5F32AI157551 (LJB), 5R35GM148258 (WDT), and DA047858 (AGH), and by the Gill Institute for Neuroscience (LJB, AGH, WDT)

## Author contributions

Conceptualization by LJB, WDT. Methodology by LJB, LCAF, JCT, AGH, WDT. Investigation by LJB, LCAF, JCT, SS, WDT. Data analysis by LJB, LCAF, JCT, SS, AGH, WDT. Visualization by LJB, LCAF. Supervision by AGH, WDT. Writing of the original draft by LJB, WDT. Review and editing of the manuscript by LJB, LCAF, AGH, WDT.

## Competing interests

Authors declare no competing interests.

## Methods

### Fly strains and husbandry

*Drosophila* stocks were raised on standard Bloomington cornmeal fly food medium at 25°C and 70% humidity on a 12/12 light/dark cycle. The following fly strains were used: *w; ppk1*.*9-Gal4* [51], *w; UAS-GCaMP6f* [52], *w; UAS-dicer2* (VDRC strain 60009), *w; nompC-Gal4* [53], and *w; 2-21-Gal4* [54] were used to build *w; ppk1*.*9-Gal4 UAS-GCaMP6f, w; ppk1*.*9-Gal4 UAS-GCaMP6f; UAS-dicer2/ K87, w; nompC-Gal4; UAS-GCaMP6f/ K87*, and *w; UAS-GCaMP6f; 2-21-Gal4* [55]. RNAi lines were obtained from the VDRC *yw; P*{*KK104185*} VDRC strain v108683 (“*UAS-ppk RNAi”*), and as a control *yw; P*{*attP* VDRC strain v60100 (“*empty attP”*) [56], and from the BDSC *yv; P*{*TRiP*.*JF01843*}*attP2* (“*UAS-bba RNAi”*), and as a control *yv; P*{*CaryP*}*attp2* (“*empty attP2”*) [57]. *w; UAS-ppk* [51] and *w; UAS-bba::mCherry* [42] were recombined to create *w; UAS-ppk UAS-bba::mCherry/TM6b Tb. w*^*1118*^ was used as the genetic control background.

### Rodents

Procedures performed with rodent species followed Institutional Animal Care and Use Committee approval at Indiana University Bloomington. Male C57BL/6J mice (Jackson Laboratories) 12 weeks of age on arrival were group-housed. Mice were given *ad libitum* access to water and food, with 48h acclimation before experiments. Because of unknown long-term effects of venom injection, all mice were euthanized 24 hours after the experiments. Sample sizes are specified in figure legends, and experimenters were blinded to treatment conditions throughout all experiments.

### Venom collection

Female velvet ants (*Dasymutilla occidentalis*) were collected at various sites in Indiana and Kentucky. Velvet ants were housed individually in glass bottles and fed with three rayon dental plugs dipped in water, 33% sugar water, and 50% honey water. The feeding plugs were replaced 1-2 times per week. Venom was collected by holding the velvet ant with forceps and allowing it to sting multiple times into a piece of parafilm stretched over an Eppendorf tube. Venom droplets were counted under a dissecting microscope (to estimate the number of stings) and then spun down into the Eppendorf tube by centrifugation at 10,000 g for 2 minutes. Venom samples were stored at -80 °C, either undiluted or diluted in buffer NEB3.1 (100 mM NaCl, 50 mM Tris-HCl, 10 mM MgCl_2_, 100 μg/mL BSA, pH 7.8).

Because of the extremely small and variable volumes of venom droplets, measuring the volume and concentration of venom samples posed a considerable challenge. Low concentration samples (“dilute venom”) were collected from individual velvet ants and contained 1-13 stings (6.2 ± 4.38 SD). These pure venom samples, which totaled much less than 1 μL, were diluted in 20 μL of buffer NEB3.1, a dilution factor of at least 1:200. Absorbance at 280 nm (A280) readings using a NanoDrop spectrophotometer can be used to estimate the concentration of protein in volumes of 1-2 μL. However, A280 depends on the presence of tryptophan and tyrosine in the proteins, which are not present in the amino acid sequence of several velvet ant venom peptides. Recognizing that these readings do not provide an accurate estimate of venom peptide concentration, we used the A280 reading to normalize how much diluted venom to use in calcium imaging experiments. 1-19 μL of diluted venom were used in a final volume of 100 μL, for a final experimental dilution factor of at least 1:20,000 (though this may vary greatly between samples). This method of normalizing resulted in consistent responses to venom within experimental groups.

For concentrated venom samples, venoms were collected and pooled from 7-9 velvet ants and contained 53-99 stings (71 ± 24.58 SD). These pure venom pools totaled ∼0.5-1 μL and were diluted to a total of 10 μL and then split in half for serial dilutions by factors of 2 to produce the relative concentration response curves in Fig. 1d. Because the starting volume of pure venom was unknown, the relative concentrations in Fig. 1d are reported by the serial dilution factor from highest concentration (1:2) to lowest concentration (1:8192). The final volume used for Ca-imaging was 70 μL, which means the highest concentration in Fig. 1d had a dilution factor of at least 1:140 and the lowest concentration of at least 1:573,440 relative to the pure collected venom.

For mouse behavior experiments with freshly collected venom, venom was diluted in PBS (137 mM NaCl, 2.7 mM KCl, 10 mM Na_2_HPO_4_, 1.8 mM KH_2_PO_4_), at dilution factors of at least ∼1:4. Fresh venom samples were kept on ice until use within less than 1 hour.

### Synthetic venom peptides

The amino acid sequences of the 24 peptides found in *Dasymutilla occidentalis* venom were obtained from Jensen et al. 2021 supplementary materials [43]. These sequences represent the mature peptide after cleavage from the translated signal peptide and propeptide which are commonly present in Hymenopteran venom peptides. The peptides were chemically synthesized by Biomatik, including C-terminal amidation if present, and delivered as lyophilized powder.

Initial screening of peptide activity in larval neurons was performed with samples of >85% purity. Subsequent larval experimental datasets and mouse experiments with peptides Do6a, Do10a, and Do13a were performed with samples of >98% purity. For larval experiments, peptides were diluted in buffer NEB3.1, and for mouse experiments peptides were diluted in PBS. Peptides Do14a and Do17a were excluded from testing because of extremely poor solubility in aqueous solution.

### Confocal calcium imaging

Wandering third-instar larvae were dissected as a fillet preparation [13]. In brief, the larva was cut open lengthwise on the ventral side, all internal organs removed (including the CNS), and the cuticle was pinned open and gently stretched to expose the peripheral sensory neurons in the dorsal larval body wall. A custom bath chamber was constructed to have a coverslip bottom (No. 1.5) through which to image sensory neurons on an inverted confocal microscope. The sides of the bath chamber were covered with magnetic strips for the use of bent metal pins with which to pin open the larval fillet at sites distant from the imaged area. Larvae were dissected in a bath buffer of modified HL3 (70 mM NaCl, 5 mM KCl, 10 mM NaHCO_3_, 0.5 mM CaCl_2_, 10 mM MgCl_2_, 5 mM HEPES, 115 mM sucrose, 4.2 mM trehalose, 5% DMSO), at pH 7.2, and ∼350 mOsm. Dissections were performed in less than 10 minutes, immediately followed by live imaging. Only one dorsal cluster of sensory neurons was imaged per larva, in an abdominal segment (A4-6), which contains one class IV da neuron, two class III da neurons, and two class I da neurons. Care was taken to avoid damaging neuron clusters during dissection and segments with any visible indication of damage were not used for imaging.

Live imaging was performed on a Zeiss LSM 5 live inverted confocal microscope with a 40x N/A 1.3 oil immersion objective. Z-stack time series were acquired using the Zen 2009 software package (Zeiss) at ∼1 volume/second, with 488 nm excitation laser and collected light with band pass filter 500-525 nm. Laser power (70%), scan speed (50 FPS), pixel dwell time (21.44 ms), pinhole size (79mm), master gain (27.9), and digital zoom (0.7) were the same for every recording, with no post-hoc image adjustments. Confocal z-stacks were set to 7-10 slices to encompass the entire depth of the sensory neuron using a fast piezo lens focus drive. Baseline fluorescence was recorded for exactly 100 frames. At frame 101, trial solutions containing venom, peptides, AITC, PBS, or NEB3.1 vehicle (always diluted in modified HL3) were added, and calcium-dependent changes in GCaMP6f fluorescence were recorded for another 200 or 400 frames. Confocal z-stacks were converted to maximum intensity projections within the Zen software and then the time-series were exported as grayscale .mov movie files for further analysis.

Care was taken to test experimental and control genotypes on the same day, with at least one positive and one negative control to ensure that venom samples and peptides retained their expected activity in every experiment.

### Quantification of calcium imaging

For analysis of calcium imaging movies, fluorescence intensity was quantified with a semi-automated MATLAB R2022b (MathWorks) code that allows the user to circle a region containing the cell body of interest. The code calculates the average luminance of a circular region of interest (9 pixel radius) centered on the cell body while correcting for lateral motion. Recordings that moved in the z-direction, causing neurons to not be fully included in the z-stack, were discarded. If the fluorescence of a neighboring cell overlapped with the cell body of interest, the data from this cell was discarded. Change in fluorescence (ΔF/F) was normalized with the formula (*F*-*F*_0_)/*F*_0_, where *F* is the fluorescence and *F*_0_ is the average baseline fluorescence in a 40-frame window within the first 100 frames with a slope closest to 0. Because of variation in the frame rates between recordings, data points were interpolated onto a standardized timeline using *csaps()* with a smoothing parameter of 1 (minimal smoothing), and aligned to the treatment time at frame 101 (assigned time = 0 s on the standardized timeline). Max ΔF/F is the maximum normalized change in fluorescence between frame 100 and the end of the recording.

### Mass spectrometry analysis

Individual samples of synthetic peptides or pooled venom were incubated for 45 min at 57 °C with 10 mM Tris(2-carboxyethyl)phosphine hydrochloride to reduce cysteine residue side chains. These side chains were then alkylated with 20 mM iodoacetamide for one hour in the dark at 21 °C. The resulting solution was desalted using ZipTip pipette tips (EMD Millipore), dried down and resuspended in 0.1% formic acid. Peptides were analyzed by LC-MS on an Orbitrap Fusion Lumos equipped with an Easy NanoLC1200. Buffer A was 0.1% formic acid in water. Buffer B was 0.1% formic acid in 80% acetonitrile. Peptides were separated on a 95-minute gradient from 5% B to 31% B over 83 minutes then to 100% B over 30 seconds and held at 100% B until the end of the run. Precursor ions were measured in the Orbitrap with a resolution of 60,000. Fragment ions were measured in the Orbitrap with a resolution of 15,000. Peptides were fragmented by HCD at a relative collision energy of 30%. Peptide MS1 extracted ion chromatograms were created at a mass accuracy of 5 ppm. The concentrations of individual venom peptides in whole venom samples were determined by ratio-metric comparison of their extracted ion chromatogram abundances to those measured in the quantified synthetic standards run in the same queue.

### Mouse behavior

To assess the effect of velvet ant venom on a mammal, nocifensive behavior and mechanical sensitivity were assessed. Initial baseline paw withdrawal thresholds to mechanical stimulation were measured, followed by intraplantar injections of venom or peptide or PBS control (vehicle) into the right hind paw (ipsilateral). Freshly collected concentrated venom was injected at a volume of 1 μL using a Hamilton syringe with a 36G needle. Synthetic venom peptides (1.2 mM in PBS) were injected at a volume of 10 μL using an insulin syringe with a 36G needle. Control injections of PBS were performed in the same manner for comparison.

Nocifensive behavior, including paw shaking, licking, and paw tapping against the enclosure, was recorded over 10 minutes immediately following the intraplantar injection. These behaviors were analyzed as indicators of nociception induced by the venom or peptides. The experimenter was blinded to the experimental group in all studies and reviewed videos of the experiment using a chronometer for precise behavioral analysis [58].

Mechanical sensitivity was assessed using an electronic von Frey anesthesiometer (model Almemo 2450, IITC Life Sciences Inc,, Woodland Hills, CA). Mice were acclimated on an elevated metal mesh table for one hour before evaluation. Mechanical withdrawal thresholds were determined by applying controlled force to the intra-plantar region of the hind paw until withdrawal, with force values recorded in grams (g) [59]. Thresholds were measured in duplicate for the ipsilateral and contralateral hind paw, and the mean of these duplicates averaged across all mice in each group, representing the mechanical paw withdrawal thresholds at each time point throughout the observation interval. Measurements were taken within a 15-minute to 24-hour interval to observe behavioral sensitivity to mechanical stimulation following the manipulation.

### Praying mantis behavior

A praying mantis (*Tenodera sinensis*) female was collected from the wild in Bloomington Indiana, at a site where we also collected velvet ants, and was temporarily housed in a plastic terrarium (14 × 8 × 6 inches) with pebbles, sticks, and water-soaked rayon plugs, and fed with live crickets 1-2 times per week. For behavioral testing, a female velvet ant was dropped into the terrarium while filming with cell phone cameras from various angles. The praying mantis was allowed to attack and attempt to feed on the velvet ant without interruption, until the velvet ant was able to escape, at which point the velvet ant was promptly removed from the terrarium.

### Statistical analysis

Data visualization and statistical analyses were performed in GraphPad Prism 10 (GraphPad Software). For serial dilutions of concentrated venom and for dose-response curves of venom peptides, the data was fit with a Sigmoidal, 4PL, curve in GraphPad Prism.

All sample sizes and statistical tests used are indicated in the figure legends. Comparison of two groups with normal distributions was performed using an unpaired Student’s t-test. One-way ANOVA was used to compare nocifensive behavior in multiple groups, followed by Tukey’s multiple comparison test. Two-way ANOVA, followed by Sidak’s post hoc test (in the case of pairwise comparisons) or Tukey’s post hoc test (for multiple comparisons) were performed, as appropriate, to compare two or more groups across time.

## Supplemental Information

**Supplemental File 1**: Figures S1-S3, Table S1, and legends for supplemental Videos S1-S4

**Video S1**: Representative exemplars of sensory neuron responses to velvet ant venom. Related to Fig. 1

**Video S2**: Representative exemplars of responses in cIII da neurons with ectopic Ppk/Bba expression. Related to Fig. 2

**Video S3**: Representative exemplars of sensory neuron responses to active venom peptides. Related to Fig. 3

**Video S4**: Velvet ant escapes predation attack by a praying mantis. Related to Fig. 4

## Notes

### Competing Interest Statement

The authors have declared no competing interest.

## References

1. Schmidt JO. The sting of the wild. Baltimore: Johns Hopkins University Press; 2016.

2. Gall BG, Spivey KL, Chapman TL, Delph RJ, Brodie Jr ED, Wilson JS. The indestructible insect: Velvet ants from across the United States avoid predation by representatives from all major tetrapod clades. Ecol Evol. 2018;8: 5852–5862.

3. Vitt LJ, Cooper WE. Feeding Responses of Skinks (Eumeces laticeps) to Velvet Ants (Dasymutilla occidentalis). J Herpetol. 1988;22: 485–488.

4. Schmidt JO, Schmidt LS, Schmidt DK. The paradox of the velvet-ant (Hymenoptera, Mutillidae). J Hymenopt Res. 2021;84: 327–337.

5. Schmidt JO. Pain and Lethality Induced by Insect Stings: An Exploratory and Correlational Study. Toxins. 2019.

6. Tracey WD, Wilson RI, Laurent G, Benzer S. painless, a Drosophila gene essential for nociception. Cell. 2003;113: 261–273.

7. Hwang RY, Zhong L, Xu Y, Johnson T, Zhang F, Deisseroth K, et al. Nociceptive neurons protect Drosophila larvae from parasitoid wasps. Curr Biol. 2007/11/29. 2007;17: 2105–2116.

8. Xiang Y, Yuan Q, Vogt N, Looger LL, Jan LY, Jan YN. Light-avoidance-mediating photoreceptors tile the Drosophila larval body wall. Nature. 2010/11/10. 2010;468: 921–926.

9. Boivin J-C, Zhu J, Ohyama T. Nociception in fruit fly larvae. Front pain Res (Lausanne, Switzerland). 2023;4: 1076017.

10. Tracey Jr. WD. Nociception. Curr Biol. 2017;27: R129–R133.

11. Burrell BD. Comparative biology of pain: What invertebrates can tell us about how nociception works. J Neurophysiol. 2017;117: 1461–1473.

12. Zelle KM, Lu B, Pyfrom SC, Ben-Shahar Y. The genetic architecture of degenerin/epithelial sodium channels in Drosophila. G3 (Bethesda). 2013;3: 441–450.

13. Ramachandran P, Budnik V. Dissection of Drosophila larval body-wall muscles. Cold Spring Harb Protoc. 2010;2010: pdb.prot5469.

14. Chen TW, Wardill TJ, Sun Y, Pulver SR, Renninger SL, Baohan A, et al. Ultrasensitive fluorescent proteins for imaging neuronal activity. Nature. 2013;499: 295–300.

15. Tsubouchi A, Caldwell JC, Tracey W. Dendritic Filopodia, Ripped Pocket, NOMPC, and NMDARs Contribute to the Sense of Touch in Drosophila Larvae. Curr Biol. 2012;22: 2124–2134.

16. Yan Z, Zhang W, He Y, Gorczyca D, Xiang Y, Cheng LE, et al. Drosophila NOMPC is a mechanotransduction channel subunit for gentle-touch sensation. Nature. 2013;493: 221–225.

17. Story GM, Peier A, Reeve A, Eid SR, Mosbacher J, Hricik T, et al. ANKTM1, a TRP-like Channel Expressed in Nociceptive Neurons, Is Activated by Cold Temperatures. Cell. 2003;112: 819–829.

18. Kwan K, Allchorne A, Vollrath M, Christensen AP, Zhang D-S, Woolf C, et al. TRPA1 Contributes to Cold, Mechanical, and Chemical Nociception but Is Not Essential for Hair-Cell Transduction. Neuron. 2006;50: 277–289.

19. Macpherson LJ, Dubin AE, Evans MJ, Marr F, Schultz PG, Cravatt BF, et al. Noxious compounds activate TRPA1 ion channels through covalent modification of cysteines. Nature. 2007;445: 541–545.

20. Neely G, Keene A, Duchek P, Chang EC, Wang Q-P, Aksoy Y, et al. TrpA1 Regulates Thermal Nociception in Drosophila. PLoS One. 2011;6: e24343.

21. Kang K, Pulver SR, Panzano VC, Chang EC, Griffith LC, Theobald DL, et al. Analysis of Drosophila TRPA1 reveals an ancient origin for human chemical nociception. Nature. 2010;464: 597–600.

22. Laursen WJ, Anderson EO, Hoffstaetter LJ, Bagriantsev SN, Gracheva EO. Species-specific temperature sensitivity of TRPA1. Temperature. 2015. pp. 214–226.

23. Jordt S, Bautista D, Chuang H, McKemy D, Zygmunt P, Högestätt E, et al. Mustard oils and cannabinoids excite sensory nerve fibres through the TRP channel ANKTM1. Nature. 2004;427: 260–265.

24. Bautista DM, Movahed P, Hinman A, Axelsson HE, Sterner O, Högestätt ED, et al. Pungent products from garlic activate the sensory ion channel TRPA1. Proc Natl Acad Sci U S A. 2005;102: 12248–12252.

25. Bautista DM, Jordt S-E, Nikai T, Tsuruda PR, Read AJ, Poblete J, et al. TRPA1 mediates the inflammatory actions of environmental irritants and proalgesic agents. Cell. 2006;124: 1269–1282.

26. Zhong L, Bellemer A, Yan H, Ken H, Jessica R, Hwang RY, et al. Thermosensory and nonthermosensory isoforms of Drosophila melanogaster TRPA1 reveal heat-sensor domains of a thermoTRP Channel. Cell Rep. 2012;1: 43–55.

27. Kang K, Panzano VC, Chang EC, Ni L, Dainis AM, Jenkins AM, et al. Modulation of TRPA1 thermal sensitivity enables sensory discrimination in Drosophila. Nature. 2011;481: 76–80.

28. Gu P, Gong J, Shang Y, Wang F, Ruppell KT, Ma Z, et al. Polymodal Nociception in Drosophila Requires Alternative Splicing of TrpA1. Curr Biol. 2019;29: 3961-3973.e6.

29. Mauthner SE, Hwang RY, Lewis AH, Xiao Q, Tsubouchi A, Wang Y, et al. Balboa binds to pickpocket in vivo and is required for mechanical nociception in drosophila larvae. Curr Biol. 2014;24: 2920–5.

30. Zhong L, Hwang RY, Tracey WD. Pickpocket is a DEG/ENaC protein required for mechanical nociception in Drosophila larvae. Curr Biol. 2010/02/18. 2010;20: 429–434.

31. Guo Y, Wang Y, Wang Q, Wang Z. The role of PPK26 in Drosophila larval mechanical nociception. Cell Rep. 2014;9: 1183–1190.

32. Ben-Shahar Y. Sensory functions for degenerin/epithelial sodium channels (DEG/ENaC). Adv Genet. 2011;76: 1–26.

33. Bianchi L. DEG/ENaC Ion Channels in the Function of the Nervous System: From Worm to Man. Adv Exp Med Biol. 2021;1349: 165–192.

34. Goodman MB, Ernstrom GG, Chelur DS, O’Hagan R, Yao CA, Chalfie M. MEC-2 regulates C. elegans DEG/ENaC channels needed for mechanosensation. Nature. 2002;415: 1039–1042.

35. O’Hagan R, Chalfie M, Goodman MB. The MEC-4 DEG/ENaC channel of Caenorhabditis elegans touch receptor neurons transduces mechanical signals. Nat Neurosci. 2005;8: 43–50.

36. Árnadóttir J, O’Hagan R, Chen Y, Goodman MB, Chalfie M. The DEG/ENaC Protein MEC-10 Regulates the Transduction Channel Complex in Caenorhabditis elegans Touch Receptor Neurons. J Neurosci. 2011;31: 12695 LP – 12704.

37. Diochot S, Baron A, Rash LD, Deval E, Escoubas P, Scarzello S, et al. A new sea anemone peptide, APETx2, inhibits ASIC3, a major acid-sensitive channel in sensory neurons. EMBO J. 2004;23: 1516–1525.

38. Escoubas P, Bernard C, Lambeau G, Lazdunski M, Darbon H. Recombinant production and solution structure of PcTx1, the specific peptide inhibitor of ASIC1a proton-gated cation channels. Protein Sci. 2003;12: 1332–1343.

39. Bohlen CJ, Chesler AT, Sharif-Naeini R, Medzihradszky KF, Zhou S, King D, et al. A heteromeric Texas coral snake toxin targets acid-sensing ion channels to produce pain. Nature. 2011;479: 410–414.

40. Diochot S, Baron A, Salinas M, Douguet D, Scarzello S, Dabert-Gay A-S, et al. Black mamba venom peptides target acid-sensing ion channels to abolish pain. Nature. 2012;490: 552–555.

41. Verkest C, Salinas M, Diochot S, Deval E, Lingueglia E, Baron A. Mechanisms of Action of the Peptide Toxins Targeting Human and Rodent Acid-Sensing Ion Channels and Relevance to Their In Vivo Analgesic Effects. Toxins (Basel). 2022;14.

42. Gorczyca DA, Younger S, Meltzer S, Kim SE, Cheng LE, Song W, et al. Identification of Ppk26, a DEG/ENaC Channel Functioning with Ppk1 in a Mutually Dependent Manner to Guide Locomotion Behavior in Drosophila. Cell Rep. 2014;9 4: 1446–1458.

43. Jensen T, Walker AA, Nguyen SH, Jin A-H, Deuis JR, Vetter I, et al. Venom chemistry underlying the painful stings of velvet ants (Hymenoptera: Mutillidae). Cell Mol Life Sci. 2021;78: 5163–5177.

44. Latorre-Estivalis JM, Almeida FC, Pontes G, Dopazo H, Barrozo RB, Lorenzo MG. Evolution of the Insect PPK Gene Family. Genome Biol Evol. 2021;13.

45. Ylla G, Nakamura T, Itoh T, Kajitani R, Toyoda A, Tomonari S, et al. Insights into the genomic evolution of insects from cricket genomes. Commun Biol. 2021;4: 733.

46. Goldberg JK, Godfrey RK, Barrett M. A long-read draft assembly of the Chinese mantis (Mantodea: Mantidae: Tenodera sinensis) genome reveals patterns of ion channel gain and loss across Arthropoda. G3 Genes|Genomes|Genetics. 2024; jkae062.

47. Svenson GJ, Hardy NB, Cahill Wightman HM, Wieland F. Of flowers and twigs: phylogenetic revision of the plant-mimicking praying mantises (Mantodea: Empusidae and Hymenopodidae) with a new suprageneric classification. Syst Entomol. 2015;40: 789–834.

48. Ronchetti F, Polidori C. A sting affair: A global quantitative exploration of bee, wasp and ant hosts of velvet ants. PLoS One. 2020;15: e0238888.

49. Gibbons M, Crump A, Barrett M, Sarlak S, Birch J, Chittka L. Chapter Three - Can insects feel pain? A review of the neural and behavioural evidence. In: Jurenka RBT-A in IP, editor. Academic Press; 2022. pp. 155–229.

50. Key B, Zalucki O, Brown DJ. Neural Design Principles for Subjective Experience: Implications for Insects. Front Behav Neurosci. 2021;15: 658037.

51. Ainsley JA, Pettus J, Bosenko D, Gerstein C, Zinkevich NC, Anderson MG, et al. Enhanced Locomotion Caused by Loss of the Drosophila DEG/ENaC Protein Pickpocket1. Curr Biol. 2003;13: 1557–1563.

52. Kim D. GCaMP6 constructs and insertions from Douglas Kim. 2013.

53. Petersen LK, Stowers RS. A Gateway MultiSite Recombination Cloning Toolkit. PLoS One. 2011;6: e24531.

54. Grueber WB, Jan LY, Jan YN. Different levels of the homeodomain protein cut regulate distinct dendrite branching patterns of Drosophila multidendritic neurons. Cell. 2003;112: 805–818.

55. He L, Gulyanon S, Mihovilovic Skanata M, Karagyozov D, Heckscher ES, Krieg M, et al. Direction Selectivity in Drosophila Proprioceptors Requires the Mechanosensory Channel Tmc. Curr Biol. 2019;29: 945-956.e3.

56. Dietzl G, Chen D, Schnorrer F, Su K-C, Barinova Y, Fellner M, et al. A genome-wide transgenic RNAi library for conditional gene inactivation in Drosophila. Nature. 2007;448: 151–156.

57. Ni JQ, Liu LP, Binari R, Hardy R, Shim HS, Cavallaro A, et al. A Drosophila Resource of 1 Transgenic RNAi Lines for Neurogenetics. Genetics. 2009;182: 1089–1100.

58. Finol-Urdaneta RK, Ziegman R, Dekan Z, McArthur JR, Heitmann S, Luna-Ramirez K, et al. Multitarget nociceptor sensitization by a promiscuous peptide from the venom of the King Baboon spider. Proc Natl Acad Sci U S A. 2022;119.

59. Tena B, Escobar B, Arguis MJ, Cantero C, Rios J, Gomar C. Reproducibility of Electronic Von Frey and Von Frey monofilaments testing. Clin J Pain. 2012;28: 318–323.

