## Supplemental File1 for "Multiple mechanisms of action of an extremely painful venom"

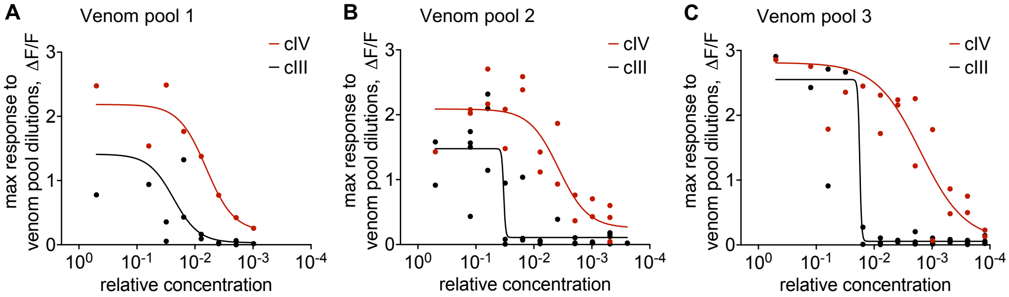


**Figure S1. Velvet ant venom has a potent nociceptor-specific component. Related to Fig. 1**

**(A-C)** Dose-response to serial dilutions of individual pooled venom samples in cIII da and cIV da neurons (*w; ppk1.9-Gal4 UAS-GCaMP6f*). Concentrated venom samples can activate both neuron types, but as the venom becomes more diluted, cIII da neurons no longer respond while the more potent cIV da-specific response remains. Data points for each neuron type are fit with a Sigmoid curve.

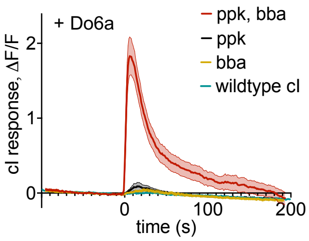


**Figure S2. Do6a activates Pickpocket/Balboa channels when co-expressed in cI da neurons. Related to Fig. 3**

Calcium imaging of Do6a (20 µM) application at time 0 s. Wildtype cI da neurons (*w; 2-21-Gal4/+; UAS-GCaMP6f/+*) do not respond to Do6a. But co-expression of *pickpocket* and *balboa* in cI da neurons (*w; 2-21-Gal4/+; UAS-GCaMP6f/UAS-ppk UAS-bba-mCherry*) renders them responsive to Do6a, while expression of *pickpocket* alone (*w; 2-21-Gal4/+; UAS-GCaMP6f/UAS-ppk*) or *balboa* alone (*w; 2-21-Gal4/+; UAS-GCaMP6f/UAS-bba-mCherry*) does not. Data represented as mean ± SEM, n = 12-16 neurons.

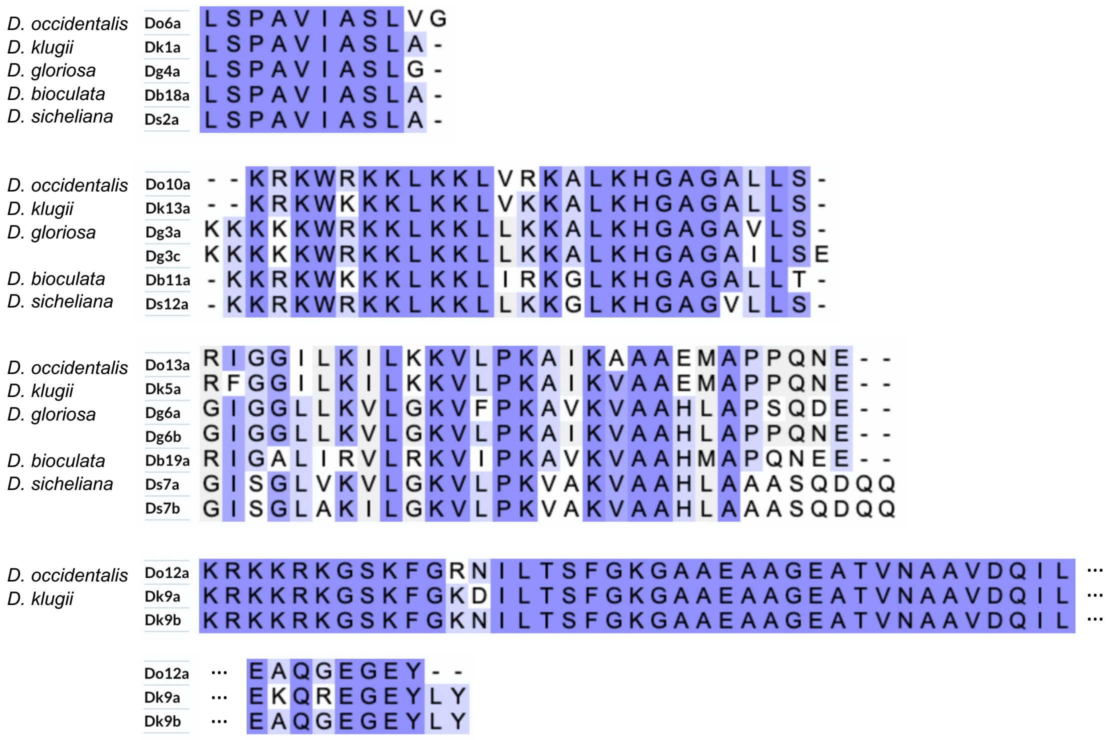


**Figure S3. Venom peptides from other velvet ant species with homology to the neuron-activating peptides of *D. occidentalis.* Related to Fig. 4**

Venom peptides with >50% amino acid identity between velvet ant species are shown here. Peptides closely related to Do6a, Do10a, and Do13a were found in all five velvet ant species whose venom proteomes have been sequenced [43], while Do12a only had homologs in two species. The color scheme represents the degree of identity of residues across the alignments. Alignments were performed in the UniProt alignment tool using Clustal Omega.

### Table S1

| Peptide name | Synthetic peptide sequence [43] | Venom sample 1 | Venom sample 2 |
|---|---|---|---|
| | | pmol per sting | pmol per sting |
| **Do6a** | **LSPAVIASLVG*** | **540** | **400** |
| Do11a | SKPSICKLVPFIPLCR* | 180 | 3.7 |
| Do5a | VFDIPNICKKRPHILRCR* | 180 | 3.7 |
| Do3a | VFTKPDICKVKPRFPPCR* | 140 | 8.2 |
| Do4a | AIYNICKIKPHLPRCR* | 120 | 3.2 |
| Do1a | DADTVAALKELIELAKKGLERI* | 110 | 280 |
| Do7a | LIFEPGICKVKPRLPLCR* | 82 | 4.2 |
| Do11c | SKPSICKLMPHIPLCR* | 59 | 0.77 |
| Do11b | SKPSICKLVPYIPLCR* | 41 | 0.73 |
| **Do13a** | **RIGGILKILKKVLPKAIKAAAEMAPPQNE** | **16** | **19** |
| Do3d | VFFTKPDICKVKPLFRPCR* | 12 | 0.18 |
| Do16a | RRRGFGNENLDKIYGGVINPQSPGVDVGKLPDDIFIQQARREFAEALAREKAERAR* | 3.8 | 1.9 |
| **Do10a** | **KRKWRKKLKKLVRKALKHGAGALLS*** | **2.9** | **1.2** |
| Do18a | RRAPPLPY | 1.1 | 0.35 |
| Do15a | LVGAALGAVELGELIHHLIKK* | 0.87 | 0.83 |
| **Do12a** | **KRKKRKGSKFGRNILTSFGKGAAEAAGEATVNAAVDQILEAQGEGEY** | **0.0040** | **0.068** |
| Do2a | VFVLPDICNERPHIPQCR* | detected in venom but not detected in standard | 73 |
| Do3b | VFTKPNICKVKPRFPPCR* | detected in venom but not detected in standard | 3.1 |
| Do3c | VFTKPDICKLRPLLRPCW* | not detected in venom sample | 0.0083 |
| Do11d | SKPTICKLVPHIPLCR* | not detected in venom sample | 0.21 |
| Do8a | SELRICRTKPRSPLCR* | not detected in venom sample | not detected in venom sample |
| Do9a | SIYNICKIKPRLPRCRGRMIGKIF | not detected in venom sample | not detected in venom sample |
| Do14a | KRGRSRGDGKKGKKPKDKIASKIGDIIKNSLNKFGVVAGETIVEKTVEAVKDALQSGEASAEEEAPSE | not detected in venom and excluded from standard | not detected in venom and excluded from standard |
| Do17a | HLGDVITDLVNKALNSL* | detected in venom but excluded from standard | detected in venom but excluded from standard |
| | | | *C-terminal amidation |

**Table S1. Peptide concentrations in two pooled venom samples**

Concentrations were determined by LC-MS by comparison to peptide standards of known concentration and were normalized to the number of stings in each pooled venom sample (126 stings in Sample 1, 58 stings in Sample 2). Peptides are presented in order of decreasing concentration based on Sample 1. Neuron-activating peptides are in bold. Do6a was the most abundant peptide in both samples.

**Video S1. Representative exemplars of sensory neuron responses to velvet ant venom. Related to Fig. 1**

Calcium imaging time series recordings are pseudocolored for fluorescence intensity. Arrows indicate cIII da cell bodies, and empty arrows indicate cIV da cell bodies. **A**, Diluted venom applied to cIV da (Fig. 1C). **B**, Diluted venom applied to cIII da (Fig. 1C). **C**, Vehicle negative control applied to cIV da (Fig. 1C). **D**, Pooled venom diluted by 1:16 applied to cIII da and cIV da (Fig. 1D). **E**, Pooled venom diluted by 1:512 applied to cIII da and cIV da (Fig. 1D). **F**, AITC applied to cIII da (Fig. 1E). **G**, AITC applied to cIII da expressing dTRPA1-C (Fig. 1E). **H**, AITC applied to cIII da expressing dTRPA1-D (Fig. 1E). **I**, Venom applied to cIII da expressing dTRPA1-C (Fig. 1F). **J**, Venom applied to cIII da expressing dTRPA1-D (Fig. 1F). **K**, AITC applied to cIV da with empty attP control for RNAi (Fig. 1G). **L**, AITC applied to cIV da with empty attP2 control for RNAi (Fig. 1G). **M**, AITC applied to cIV da with ppk RNAi (Fig. 1G). **N**, AITC applied to cIV da with bba RNAi (Fig. 1G). **O**, Venom applied to cIV da with empty attP control for RNAi (Fig. 1H). **P**, Venom applied to cIV da with empty attP2 control for RNAi (Fig. 1H). **Q**, Venom applied to cIV da with ppk RNAi (Fig. 1H). **R**, Venom applied to cIV da with bba RNAi (Fig. 1H). Video frame rate is sped up 10 times.

**Video S2. Representative exemplars of responses in cIII da neurons with ectopic Ppk/Bba expression. Related to Fig. 2**

Calcium imaging time series recordings are pseudocolored for fluorescence intensity. Arrows indicate cIII da cell bodies. **A**, Venom applied to cIII da expressing Ppk alone (Fig. 2A-B). **B**, Venom applied to cIII da expressing Bba alone (Fig. 2C-D). **C**, Venom applied to cIII da co-expressing Ppk and Bba (Fig. 2E-F). **D**, Vehicle negative control applied to cIII da co-expressing Ppk and Bba (Fig. 2G-H). Video frame rate is sped up 10 times.

**Video S3. Representative exemplars of sensory neuron responses to active venom peptides. Related to Fig. 3**

Calcium imaging time series recordings are pseudocolored for fluorescence intensity. Arrows indicate cIII da cell bodies, and empty arrows indicate cIV da cell bodies. **A**, Do6a (100μM) applied to cIII da and cIV da (Fig. 3A-B). **B**, Do10a (100μM) applied to cIII and cIV da (Fig. 3A-B). **C**, Do12a (100μM) applied to cIII and cIV da (Fig. 3A-B). **D**, Do13a (100μM) applied to cIII and cIV da (Fig. 3A-B). **E**, Do6a applied to cIV da with empty attP control for RNAi (Fig. 3C). **F**, Do6a applied to cIV da with empty attP2 control for RNAi (Fig. 3C). **G**, Do6a applied to cIV da with ppk RNAi (Fig. 3C). **H**, Do6a applied to cIV da with bba RNAi (Fig. 3C). **I**, Do6a applied to cIV da (Fig. 3D). **J**, Do6a applied to cIII da (Fig. 3D). **K**, Do6a applied to cIII da co-expressing Ppk and Bba (Fig. 3D). Video frame rate is sped up 10 times.

**Video S4. Velvet ant escapes predation attack by a praying mantis. Related to Fig. 4**

A praying mantis (female *Tenodera sinensis*) is exposed to a Scarlet Velvet Ant (female *Dasymutilla occidentalis*). The praying mantis captures the velvet ant and attempts to consume it. The velvet ant's hard carapace prevents immediate injury and allows the velvet ant to defend itself by stinging the mantis. A visible sting occurs on the mantis foreleg femur/trochanter, and the mantis immediately startles and releases the velvet ant from its grip. In 5 out of 5 similar trials, the velvet ant successfully escaped from predation by the praying mantis.
